# Uncovering the Morphological Evolution of Language-Relevant Brain Areas

**DOI:** 10.1101/2023.03.17.533103

**Authors:** Guillermo Gallardo, Cornelius Eichner, Chet C. Sherwood, William D. Hopkins, Alfred Anwander, Angela D. Friederici

**Affiliations:** Department of Neuropsychology, Max Planck Institute for Human Cognitive and Brain Sciences, Leipzig, Germany; Department of Anthropology, The George Washington University, Washington DC, USA; Department of Comparative Medicine, Michale E. Keeling Center for Comparative Medicine, The University of Texas MD Anderson Cancer Center, Bastrop, Texas, USA

**Author notes:** Equal Contribution (Guillermo Gallardo, Cornelius Eichner). **Corresponding Author** Guillermo Gallardo, **E-Mail**.

## Abstract

Human language is supported by a cortical network involving Broca’s area which comprises Brodmann Areas 44 and 45 (BA44, BA45). While cytoarchitectonic homolog areas have been identified in nonhuman primates, it remains unknown how these regions evolved to support human language. Here, we use histological data and advanced cortical registration methods to precisely compare the morphology of BA44 and 45 between humans and chimpanzees. We found a general expansion of Broca’s areas in humans, with the left BA44 enlarging the most, growing anteriorly into a region known to process syntax. Together with recent functional studies, our findings show that BA44 evolved from a purely action-related region to a more expanded region in humans, with a posterior portion supporting action and an anterior portion supporting syntactic processes. Furthermore, our findings provide a solution for the longstanding debate concerning the structural and functional evolution of Broca’s area and its role in action and language.

## Introduction

Language processing is a human trait that involves Broca’s area in the inferior frontal gyrus. Previous studies suggest an involvement of this area in the perception, execution, and imitation of actions [1]. Moreover, homologous areas in nonhuman primates have similarly been shown to be involved in actions of the orofacial muscles and upper limbs [2]. However, the relationship between action and language, and how Broca’s area evolved to support them remains elusive. A longstanding debate persists regarding the relationship between language and action, and whether they share the same neural basis. Within this debate, there are currently two opposing views. The first view states that language emerged from action expressed in communicative gestures, and thus they share a common basis [2–4]. The second view sees language as a cognitive ability independent of action [5,6].

Both views - favoring and opposing a shared basis for language and action - built their arguments on theoretical and empirical grounds [2–4]. At the theoretical level, the debate focuses on the (di)similarity between the hierarchical structure of linguistic syntax, as a core aspect of human language, and that of goal-directed sequential actions. While some argue that action can be described using a hierarchical structure of subgoals [7,8], others claim that such a description does not meet the definition of hierarchy in language [5]. In this way, the theoretical arguments are stalled. Meanwhile, at the empirical level, several studies in humans have found action to recruit Broca’s area [9,10], an area primarily related to language, thus suggesting a functional co-dependence between action and language [1,2,11]. However, these studies did not compare actions versus language, nor tried to identify their exact location, thus making it hard to understand if the same regions activate for both processes.

To date, there are only two studies directly comparing the neural underpinning of action and language in humans. One is a meta-analysis comparing peak activations of syntactic tasks against motor-related ones [12], and the other is a functional imaging study comparing tool use with language in a within-subject design [13]. Both of these studies found that language and action recruit non-overlapping areas of Broca’s area, with language being processed more anterior than action. Moreover, the meta-analysis shows that Brodmann Area 44 (BA44), the cytoarchitectonic defined posterior division of Broca’s area [14,15], is functionally divided in two regions. In BA44, language recruits the anterior part and action recruits the posterior one [12].

To help settle the debate on the language/action relationship we can turn to our close evolutionary relatives. Anthropoid primates, such as chimpanzees and macaques, possess a cytoarchitectonically similar Broca’s area homolog that, as in humans, functionally responds to action [10]. Moreover, there is evidence that great apes can master some aspects of language using augmentative or alternative communication systems such as sign language or visual graphic symbols [16]. However, only humans possess the faculty of language when defined by the development of syntax and the ability to create multiword utterances following a set of grammatical rules. Hence, a cross-species comparison between the linguistic human brain and one of our closest living relatives, namely chimpanzees, may shed light on the neural basis of action and language.

Earlier cross-species comparisons have shown that the prefrontal cortex is a region that allometrically scales to increase at a disproportionate rate across primates, leading to a relatively large size in the human brain [17,18]. Particularly, the cytoarchitectural regions BA44 and BA45 were found to be up to 6.6-fold larger in humans than in chimpanzees (1.3-fold and 1.4-fold larger than expected, respectively, after correcting for overall cortical enlargement) [19]. Furthermore, based on histological studies, it has been shown that Broca’s subregions BA44 and BA45 differ between humans and chimpanzees in terms of their asymmetry. Human BA45 reaches its leftward volumetric asymmetry by the age of 5 years during development. Human BA44 only reaches its asymmetry by the age of 11 years [20] when children acquire full proficiency in semantic and syntactic knowledge [21]. In contrast, in chimpanzees, neither BA44 nor BA45 develop volumetric asymmetry [19].

Here, we examined the evolution of Broca’s area by comparing cytoarchitectural segmentations of BA44 and BA45 in humans and chimpanzees, derived from published histological data. Leveraging advanced cortical registration methods [22,23], we aligned the brains of chimpanzee and human, enabling us to perform a direct comparison of the segmentations across species. Our analysis confirms that Broca’s area expanded in humans, with left BA44 being the subregion that enlarged the most. Furthermore, we show that when mapped into the human brain, the chimpanzee left BA44 only covers the posterior region related to action, while having virtually no overlap with the anterior syntax regions. Our results provide strong evidence that BA44 evolved from a pure action region, as found in our closet living ape relatives, to a bipartite system recruiting action posteriorly, and syntax anteriorly. In this way, we provide a complete picture of Broca’s area evolution and offer a solution to the longstanding debate of BA44’s role in language and action.

## Results

### Symmetry of Broca’s Area Homolog in Chimpanzee and a Surface Probabilistic Atlas

Through a semi-supervised pipeline (summarized in Fig. 1a) we precisely reconstructed the cortical surface of nine chimpanzee brains from their structural MRI data. Having their surface representation, we projected both BA44 and BA45 volumetric histological segmentations [19] to each individual surface. We examined all individuals for evidence of surface area asymmetry of both histologically defined regions (BA44 and BA45) using a Wilcoxon signed-ranks test. Although there was considerable asymmetry in some individuals (see Fig. 1b) both BA44 and BA45 showed no asymmetry at the population level (BA44 T=16, p=0.49; BA45 T=10, p=0.16.

**Figure 1.**
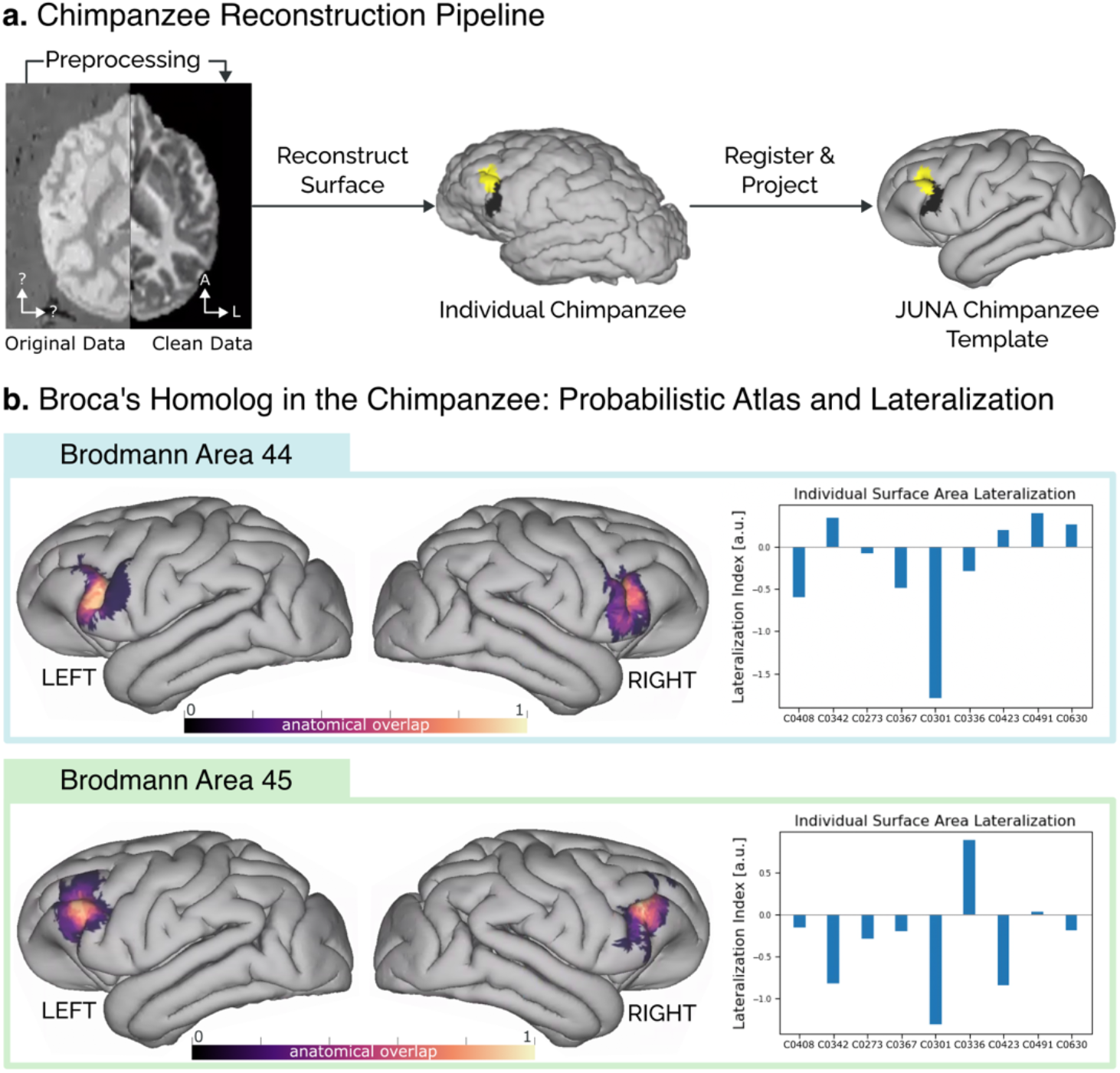
**(A)** Reconstruction pipeline for the cytoarchitectonic surface maps. First, the raw MRI data were cleaned using noise reduction and contrast inversion. Next, the individual surfaces were reconstructed in FreeSurfer. The individual maps of BA44 and BA45 are displayed in black and yellow, respectively. Finally, the individual surfaces and cytoarchitectural maps were registered to the JUNA template surface **(B)** Probabilistic atlas of regions BA44 and BA45 in the chimpanzee brain, derived from the individual maps, alongside the lateralization index for each individual brain.

To enable comparison across subjects, we co-registered the individual brain surfaces to the surface reconstruction of the JUNA [24] chimpanzee template (see Fig. 1a). On the JUNA surface we averaged all the individual segmentations, deriving a high-quality probabilistic atlas of BA44 and BA45 homologs in the chimpanzee brain (Fig 1b). The resulting atlas is open access and available for direct download (see Data and Code Availability Statement).

### Comparison Between Human Broca’s Area and its Chimpanzee Homolog

Leveraging advanced surface registration [22,23], we co-registered the JUNA surface to the surface reconstruction of the MNI-2009c human template (Fig. 2a) [25]. This enabled us to compare the human BA44 and BA45 histological atlases derived by Amunts et al. [15], with our probabilistic atlas of the chimpanzee homolog (Fig. 2b).

**Figure 2.**
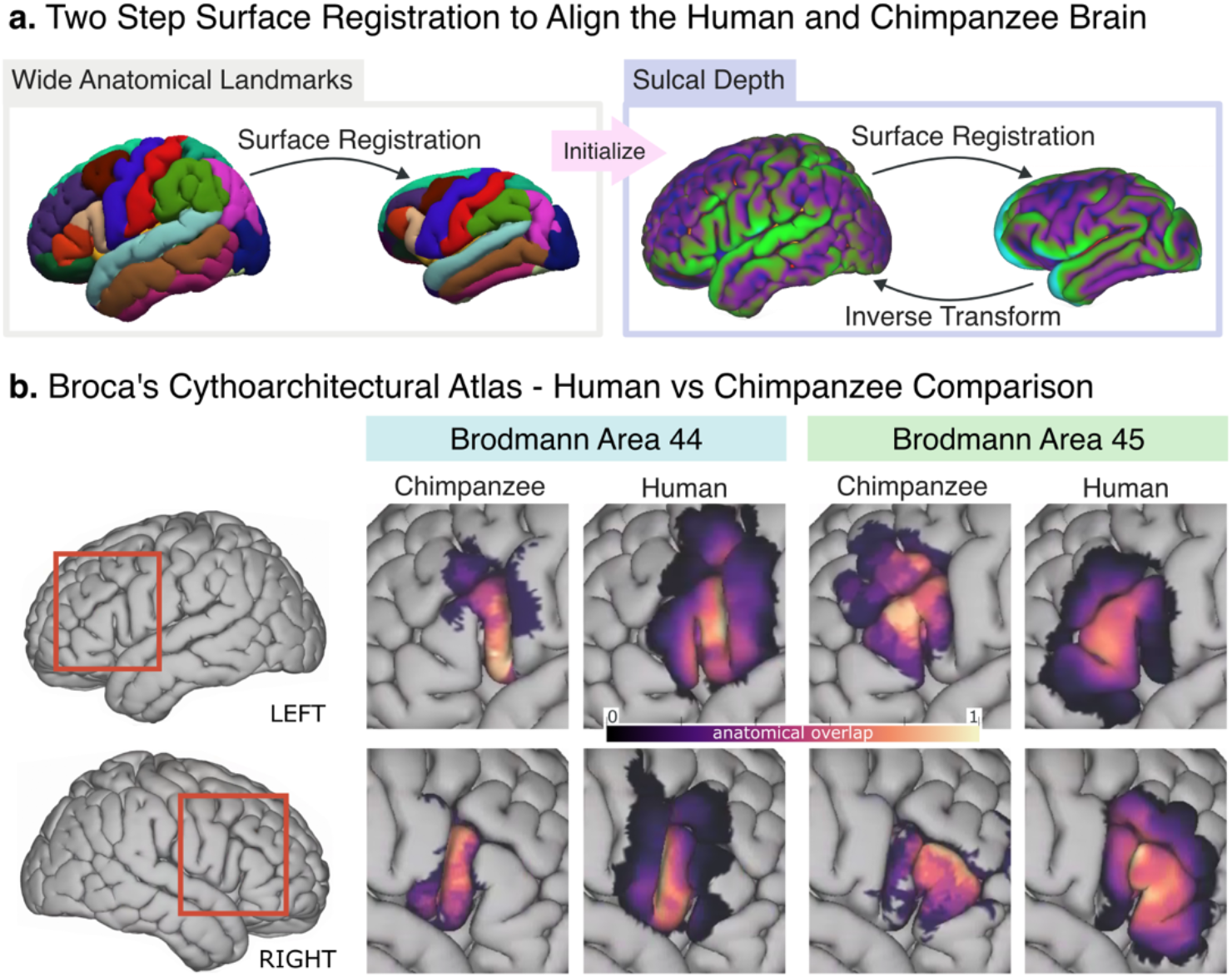
**(A)** Two-step surface registration, in the first step we align gross anatomical landmarks. This first alignment is then used to start a more granular one, based on sulcal depth. **(B)** Side-by-side comparison of our chimpanzee probabilistic atlas with the human population overlap of Amunts et al. [15] in the human brain template. Left BA44 is the area that grew the most and shows a large anterior expansion which is not present in Right BA44.

After projecting the chimpanzee segmentations to the human brain, we computed their volumes using the MNI template’s cortical thickness. We found the chimpanzee BA44 to have an average size of 2714 mm^3^ (SD: 1059) in the left hemisphere, and 2179 mm^3^ (SD: 1184) in the right hemisphere. In contrast, Amunts et al. [15] reported the human BA44 to have an average size of 3839 mm^3^ (SD: 2277) in the left hemisphere, and 2527 mm^3^ (SD: 1597) in the right hemisphere. This means that, when scaled and projected to a same surface template, the human BA44 is 1.42 times larger in the left hemisphere than in the chimpanzee, and 1.16 times larger in the right hemisphere. Moreover, Figure 2b shows that such enlargement is likely the result of a large anterior expansion, not present in the right BA44.

For the chimpanzee BA45, the average size after projecting to the human brain (Fig. 2b) was 3168 mm^3^ (SD: 1212) and 2347 mm^3^ (SD: 1435) for the left and right hemispheres, respectively. For the same region in humans, Amunts et al. [15] reported an average size of 3242 mm^3^ (SD: 1149) and 3173 mm^3^ (SD: 1637) for the left and right hemispheres, respectively [15]. In comparison, the human BA45 was only 1.02 times larger in the left hemisphere than the chimpanzee’s homolog area, while being 1.35 times larger in the right hemisphere.

### Comparing the Projection of Chimpanzee BA44 with Human Functional Maps

To better understand the behavioral role of the observed expansion, we computed the overlap of the projected chimpanzee BA44 with sub-divisions of human BA44 related to action and syntax [12,26–28]. To compare only with the core chimpanzee BA44, we thresholded the projected atlas at the 0.5 level. We found that the chimpanzee BA44 overlapped most with the regions involved in action [12,26] (Fig. 3 left, Table 1, and Sup. Fig. 1). The highest overlap was found with the area Clos 4, associated with action imagination [26], of which 34% was contained by the chimpanzee’s BA44. Following this were the regions Clos 1 (26% contained, associated with phonology and overt speech tasks [26]), Clos 5 (20%, associated with phonology and semantics [26]), Papitto’s region (18%, associated with action execution/imitation [12]), and Clos 2 (7%, associated with semantics, orthography, and covert speech [26]). In contrast, the region Clos 3, associated with basic syntactic operations [26–28], had only a 3% overlap with the chimpanzee BA44. Similar results were obtained when comparing across different levels of thresholding (see Sup. Fig. 1).

**Figure 3.**
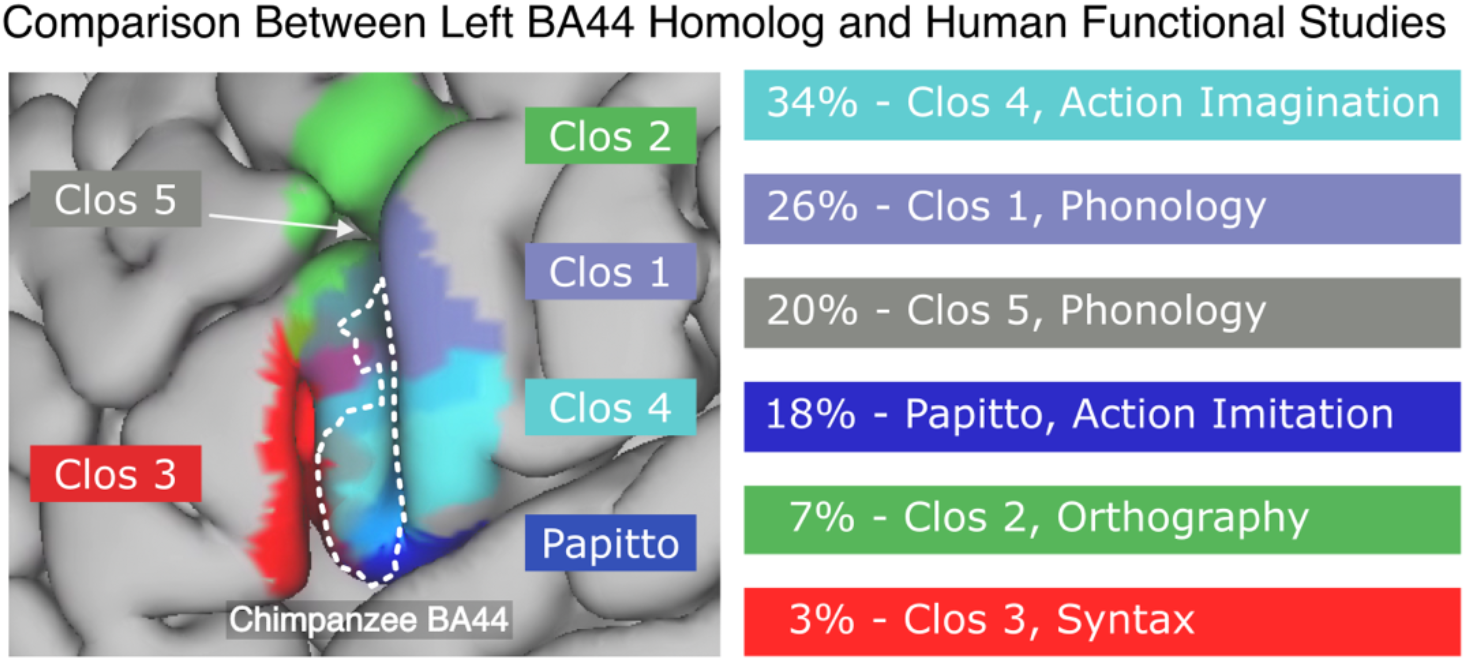
Percentage of overlap between the chimpanzee BA44 and functional subdivisions of the human BA44 [12,26–28]. Action-related regions present the highest overlap with action-related areas and virtually no overlap with the syntax area. The chimpanzee BA44 atlas was thresholded at 0.5 to maintain only its core area. The functions being reported are those with the highest P (Activation | Domain) as reported by Clos et al. [26] and Papitto et al. [12], excepting Clos 1, which was originally reported to be a syntax area, but further studies did not find to be involved in basic syntactic operations [26,27].

**Table 1.** Anatomical overlap between functional areas [12,26] with the projected BA44 chimpanzee thresholded at 0.5. The functional role of each area, as reported by their authors, is stated in the third column. For Clos regions, we report the functions with the highest P (Activation | Domain). *: It is important to notice that, even though Clos was originally reported to be a syntax area, further studies did not find it to be involved in basic syntactic operations [27,28].

| **Region** | **Area Contained by BA44** | **Associated Functions** |
| --- | --- | --- |
| Clos 4 | 34% | Action Imagination |
| Clos 1 | 26% | Phonology, Syntax* |
| Clos 5 | 20% | Phonology, Semantics |
| Papitto | 18% | Action Execution / Imitation |
| Clos 2 | 7% | Orthography, Working Memory |
| Clos 3 | 3% | Syntax, Phonology |

## Discussion

In this study, we leveraged comparable cytoarchitectonic segmentations of BA44 and 45 from chimpanzees and humans to study evolutionary changes in their size and distribution on the cortical surface. Particularly, we used advanced algorithms [22,23] to align human and chimpanzee brains, enabling a direct comparison of their (a)symmetry, size, alignment, and functional relevance.

### BA44 Became Increasingly Left Lateralized in Evolution

We tested for asymmetry in surface area for both BA44 and BA45 in chimpanzee brains. In consistency with Schenker et al. [19] volumetric analysis, we found both regions to show no statistical difference in size, thus being symmetric at the population level. This result is in clear contrast with the strongly left lateralized BA44 in humans [15]. Since both chimpanzee and human segmentation were obtained through similar histological procedures, the results support the conclusion that the asymmetry of BA44 developed after humans diverged from our last common ancestor with chimpanzees. Sample sizes are relatively small for histological analyses in both humans and chimpanzees and this conclusion should be tested further with a larger number of subjects.

### BA44 Expanded the Most in Humans Relative to Chimpanzees, Extending Anteriorly

We projected the BA44 and BA45 of each chimpanzee to the surface of a human cortical template and computed an average volume using the template’s cortical thickness. Assuming the MNI template is representative of the Amunts et al. [15] population, their reported volumes for humans are directly comparable with our scaled-up volumes for chimpanzee. Our comparison revealed that BA44 enlarged by a factor of 1.42 and 1.16 in the left and right hemispheres beyond the amount of overall cortical expansion, respectively, in this cross-species comparison. Meanwhile, BA45 enlarged only in the right hemisphere, by a factor of 1.35. Our results show that Broca’s area enlargement is remarkable in context of the evolution of human prefrontal cortex size [17,19], while showing a clear anterior expansion of left BA44, the area that enlarged the most.

### The Chimpanzee BA44 Overlaps with Human Areas Related to Action and not Syntax

We directly compared the chimpanzee BA44 atlas, projected to the human cortical surface, with subdivisions of Broca’s area derived from human functional imaging studies. We found that the core chimpanzee BA44 overlapped solely with areas related to action, with the greatest overlap found (in descending order) with regions related to action imagination, phonology, and action execution/imitation [26] [12]. Indeed, we found almost no overlap between the core BA44 chimpanzee homolog and the Broca’s subdivision involved in basic syntax operations (Clos et al. [26], Zaccarella et al. [27,28]). These results were consistent across multiple levels of thresholding for the chimpanzee BA44 probabilistic map.

### BA44 Evolved into a Bipartite Region to Support both Action and Syntax in Humans

Recent functional imaging studies found that both language and action recruit non-overlapping subdivisions of Broca’s area in the human brain, with language being processed more anteriorly than action [12,13,26–28]. Moreover, it has been found that particularly left BA44 segregates action and language processing in two distinct sub-regions, with language recruiting its anterior part and action the posterior one [12,26–28].

In this study, we have found strong evidence that the left human BA44 evolved to accommodate syntax through an anterior expansion of action-related regions in the inferior frontal cortex. Particularly, we have shown that this region enlarged more than the rest of Broca’s area subdivisions, and that, when mapped into the human brain, its chimpanzee homolog only corresponded to cortical regions known to support action. Hence, our results support the theory of a bipartite left BA44, with an anterior segment serving syntax and a distinct posterior segment serving action. Further, we have shown that this bipartite nature is likely the result of an evolutionary process. A process in which the pure action precursor of left BA44 expanded anteriorly to give rise to the core component of our human language – syntax.

## Materials and Methods

### Cytoarchitecture Segmentation of Broca’s Area in Human Brains

We downloaded the publicly available data from Amunts et al. [15], in which the left and right BA44 and BA45 were manually segmented on 10 subjects following histological procedures. While the individual maps are not available, the Julich institute has released the probabilistic cytoarchitectural map of both areas derived from the Amunts’ dataset.

### Cytoarchitecture Segmentation of Broca’s Area Homolog in Chimpanzee Brains

Our chimpanzee cytoarchitectural data comes from a previous study, in which both BA44 and BA45 were bilaterally delineated and guided by the same cytoarchitectonic criteria defined for humans by Amunts et al. [19]. Whole-brain MRI data were acquired ex-vivo for all chimpanzees (Supplementary Information). From the population of 12 chimpanzees, we discarded 3 based on inadequate corresponding MRI data quality, retaining 9 subjects (*Pan troglodytes*, 5/4 males/females, age = 32.8 ± 11.8 years, age range = 12–44.5 years). The demographics of the included chimpanzees are summarized in Supplementary Table S1. For additional information on the data acquisition please refer to the original publication [19].

### Broca’s Homolog in the Chimpanzee: Deriving a Probabilistic Atlas and Studying Population Symmetry

We derived a probabilistic atlas of Broca’s area homolog from the individual cytoarchitectonic segmentations of BA44 and BA45, and their associated ex-vivo MRI scans. We performed all the analysis on the reconstructed cortical surfaces, as surface analysis better captures and aligns brains based on their gyrification, thus being more robust than volumetric analysis [29].

The procedure can be summarized in five steps: (I) reconstruct 3D brain surfaces from the ex-vivo MRI scans using FreeSurfer, (II) project each chimpanzee’s BA44 and BA45 volumetric segmentation to their corresponding surfaces, (III) register all surfaces to a common template, namely the JUNA chimpanzee brain template [24], (IV) map the individual cytoarchitectural regions to the JUNA template, and (V) aggregate them to derive a high-quality probabilistic atlas. See Fig. 1a. for a graphical explanation, and the Supplementary Methods for a detailed explanation of each step. The processing scripts for the computation of the open access atlas are readily available for download (see Data and Code Availability Statement).

We further leveraged the individual reconstructions to study the surface-area asymmetry of Broca’s homolog in the chimpanzee brain. For this, we computed the areas of BA44 and BA45 on each individual chimpanzee, and tested their bilateral symmetry through a Wilcoxon sign-ranks test.

### Mapping Chimpanzee Cytoarchitectural Maps to the Human Brain

To enable cross-species comparison, we aligned the cortical reconstruction of JUNA template [24] to that of the human MNI template (ICBM152 9c Asymmetric) [25]. Given the differences in brain shape and volume, we opted to use surface-based registration algorithms, which have been proven successful in aligning the brains of chimpanzees and humans [23].

Based on the work of Eichert et al. [23] we performed the surface-based registration in two stages. In the first stage, we performed a first alignment of the brain templates using gross anatomical regions. Specifically, we aligned the brains based on their inferior frontal gyrus, as defined by the Desikan atlas (Fig. 2a) [30]. Starting from that rough alignment, we then carried a more granular registration based on the sulcal patterns. For a detailed explanation of each stage please refer to the Supplementary Methods as well as the open access processing script (see Data and Code Availability Statement).

### Expansion of BA44 & BA45 in Humans Relative to Chimpanzees

In their histological study, Amunts et al. [15] report the average gray-matter volume for human BA44 and BA45. Since our chimpanzee regions stem from similar histological procedures, we can study how much BA44 and BA45 expanded through evolution by mapping them to a common space, and comparing their size across species.

Having morphed chimpanzee BA44 and BA45 to the human template, we computed their individual volumes using the MNI template cortical thickness. In this way, we obtained volumes for chimpanzee Broca’s area subregions that are scaled up and projected onto the template human cortical surface. Assuming the MNI template is representative of the Amunts et al. [15] population, their reported volumes for humans are directly comparable with our scaled-up volumes for chimpanzee.

### Functional Aspects of the BA44 Homolog in the Human Brain

We aimed to understand the relation between function and the location of the projected chimpanzee Broca’s area homolog - with a particular interest in language and action. For this, we projected the functional subdivisions of human BA44 defined by Papitto et al. [12] and Clos et al. [26] to the MNI cortical surfaces. There, we compared them to the core chimpanzee BA44, obtained by thresholding the atlas at the 0.5 level, i.e. the points in the surface where the majority of the chimpanzee population had their BA44 located. Particularly, for each functional region we computed their overlap with the chimpanzee BA44, defined as how much of the functional area was contained by the chimpanzee BA44.

## Supplementary Information

### Chimpanzee Data

Five chimpanzees had been wild-caught and lived in captivity since then. The remaining four chimpanzees were born and raised in captivity. All subjects lived in social groups ranging from 2 to 13 individuals at Emory National Primate Research Center and were housed according to institutional guidelines. An overview of the included sample demographics is summarized in Table S1 [19].

The brains were extracted after subjects died from causes unrelated to the study. The brains were removed within 14h of each subject chimpanzee’s death and immersion fixed in 10% formalin. All brain samples were placed in agarose gel to increase mechanical stability for the MRI acquisition [19].

**Table S1:** Demographics of the included subjects.

| Specimen | Sex | Age (y) | Rearing |
| --- | --- | --- | --- |
| C0273 | m | 40,0 | Wild |
| C0336 | f | 44,0 | Wild |
| C0342 | f | 35,4 | Wild |
| C0367 | m | 41,2 | Wild |
| C0408 | f | 44,5 | Wild |
| C0301 | m | 35,9 | Captive - Nursery |
| C0423 | m | 24,0 | Captive - Nursery |
| C0491 | m | 18,5 | Captive - Mother |
| C0630 | f | 12,0 | Captive - Mother |

### Chimpanzee MRI Data Acquisition

Whole brain MRI scans of the ex-vivo samples were acquired using a 1.5 T MRI system (GE Medical Systems, Milwaukee, WI) with software version 8.3. Anatomical T1-weighted data were coronally acquired with the following parameters: TR = 666.7 ms, TE = 14.5 ms, 80 slices, slice thickness = 1.5 mm, in-plane matrix size = 256 × 256, in-plane FoV = 160 mm × 160 mm, in-plane resolution 0.625 × 0.625 mm.

### Ex-Vivo Data Challenges

Processing ex-vivo MRI data involves specific challenges, which must be addressed to use standard in-vivo processing pipelines. First, the observed MRI contrast is affected by fixation-induced changes in relaxation times, with heavy implications on the utility of in-vivo contrast-based segmentation tools. Second, the brains are extracted and embedded in agarose with high signal intensity - making it challenging to use typically employed brain extraction tools. Third, air bubbles trapped in gyri or agarose may cause local distortions or contrast changes and need to be carefully excluded from the overall reconstruction.

### Preparing Ex-Vivo Data for FreeSurfer

We implemented a semi-automated pipeline to prepare the ex-vivo MRI of each chimpanzee to be later processed in FreeSurfer 6.0.0. In total, 5 steps were necessary: (I) Align the brains to the AC-PC convention (thus simplifying their visualization across tools), (II) separate the brain from the background using an in-house semi-supervised algorithm (to avoid errors during FreeSurfer’s brain extraction), (III) standardize the brain resolution to 0.7mm isometric (improves the memory and time needed to process each brain), (IV) invert the contrast on those chimpanzees with white matter darker than gray matter (FreeSurfer expects a T1-like contrast), and (V) apply a Gaussian blurring of 0.4mm to smooth intensity values within both gray-matter and white-matter tissue.

### Semi-supervised FreeSurfer Pipeline for Chimpanzees

Given that FreeSurfer was designed to process human brains, we had to use a configuration slightly different from the default to process the chimpanzee brains. Particularly, we: (1) Remove the brain extraction process, since we had already separated the brain from the background (see previous section), (2) run the whole pipeline using the “high resolution” flag, which stops FreeSurfer from resizing the voxels to 1 mm isometric, (3) removed the “Talairach check” since it fails when processing non-human brains, (4) manually normalized the brain intensity, such that the average white-matter value was 110, and (5) manually checked the white-matter segmentation, fixing it when necessary (though this is a normal step for processing humans as well). To simplify controlling the pipeline, each step was run separately and followed by a thoughtful visual control.

### Registering the Individual Chimpanzee Surfaces to a Common Template

To compare the chimpanzees in a common space, we registered the individual brains to the JUNA chimpanzee template. Given that the JUNA template is volumetric, we first derived its 3D brain surface reconstruction with the same FreeSurfer pipeline used on the remaining chimpanzee brains. Once the reconstruction was available, we performed surface-based registration with MSM, using the JUNA template as the source and each individual chimpanzee brain as the target. Surface-based alignment has been proven to be more robust and precise than volumetric-based registration [29].

Since the chimpanzee brains are consistent across individuals, we used the original pairwise implementation [31], which results in a fast yet accurate registration. For the exact configuration file please see the supplementary file ‘chimp-to-juna-msm-config.txt’.

### Deriving the Chimpanzee Cytoarchitectural Probabilistic Atlas

By inverting the transformations derived in the previous step, we were able to project each individual cytoarchitectural segmentation to the template. Having them in a common space, we then averaged the segmentations to derive a high-quality probabilistic map of both BA44 and BA45 in the chimpanzee brain.

### Co-Registering the Human and Chimpanzee Brain

To enable a cross-species comparison between humans and chimpanzees, we performed an alignment between the brains of both species. Specifically, our goal was to align the JUNA chimpanzee template with the human MNI template (ICBM152 9c Asymmetric). Given the difference in volume and proportion of white/gray matter between humans and chimpanzees, volumetric registration is sub-optimal. Hence, we performed a surface-based alignment which is not subject to this restriction. Surface-based registration has been shown successful to compare brains across species, due to its ability to characterize and align brains based on their gyrification [23]. Before registering the brains, we first had to process the MNI template to reconstruct its 3D brain surface representation. This was achieved using the FreeSurfer pipeline with default parameters. FreeSurfer is perfectly suited for processing human brains, for which no inconveniences were raised during the process.

To perform the surface-based registration between the JUNA template surface and the MNI one we used the strain-based regularization version of MSM [22]. This algorithm produces smoother warps and better alignment in the presence of noise, and thus should be preferred when aligning brains across species. The registration was performed in two stages. In brief, we first aligned the brains based on gross anatomical landmarks, to then improve the alignment based on local sulcal data.

For the first stage, we derive a binary mask for the Inferior Frontal Gyrus (IFG) in both templates. The IFG is in fact extracted from the anatomical parcellation produced by FreeSurfer (known as the Desikan atlas). Since humans and chimpanzees have pronounced landmarks surrounding the IFG (Sylvian Fissure, Precentral Sulcus, and Lateral Sulcus), the Desikan atlas adequately represents it in both species [32]. We further improved this first registration by aligning the templates using local sulcal data. Since the human brain is bigger and thus has higher resolution, we performed the alignment using the MNI brain as the source and the JUNA template as the target. For the exact configuration file please see the configuration file ‘human-to-chimp-msm-config.txt’

### Supplementary Figures

**Supplementary Figure 1:**
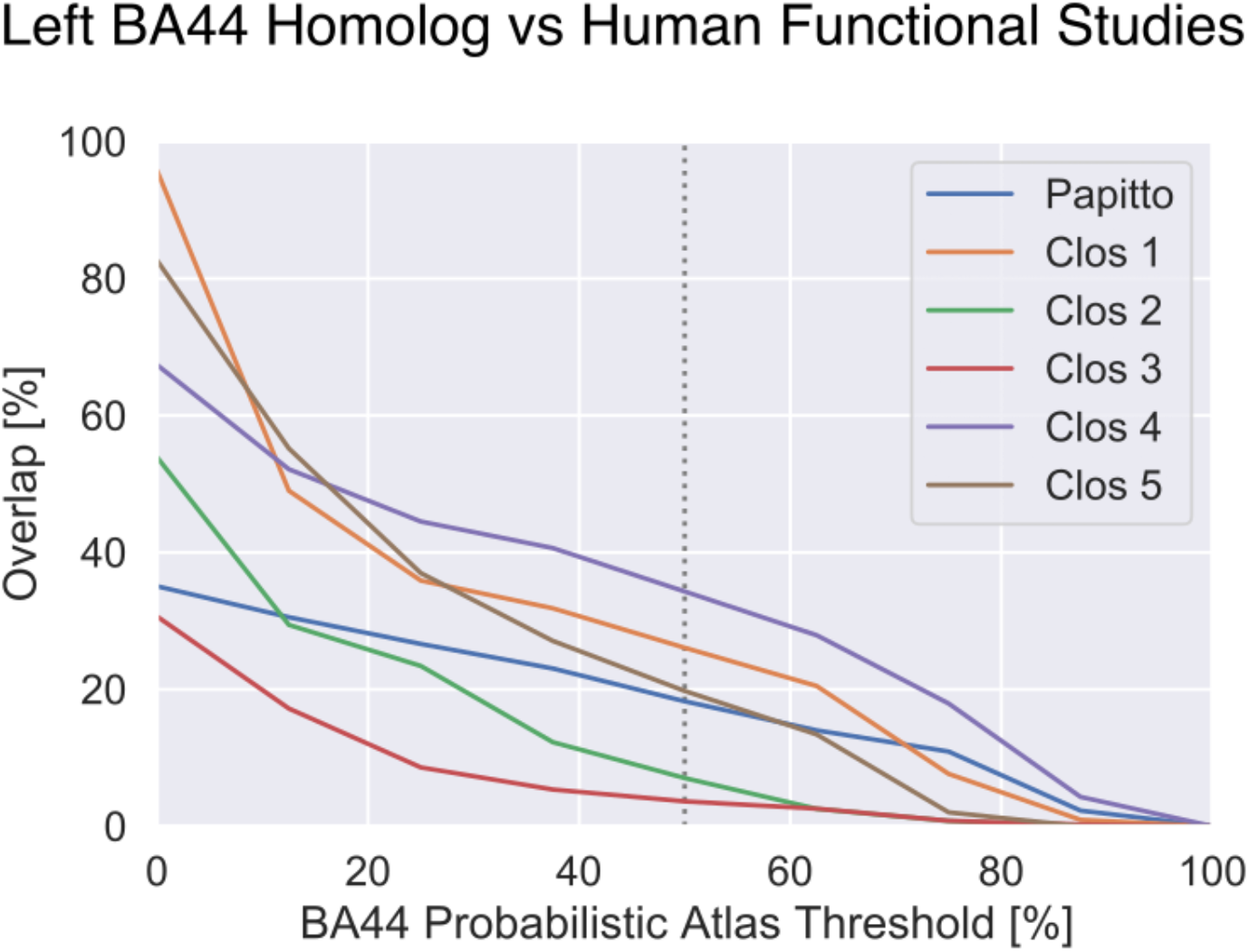
Comparing the overlap of Functional Subdivisions of the Human BA44 and the Chimpanzee BA44 Probabilistic Atlas at different levels of threshold. Notice that only the probabilistic atlas is being thresholded. As expected, the overlap decreases as the area of BA44 shrinks (the threshold increases). For all threshold levels, the least overlapping region is the syntax-related area Clos 3.

## Acknowledgment

This study was funded by the Max Planck Society under the inter-institutional funds of the president of the Max Planck Society for the project “Hominoid Brain Connectomics” (M.IF.A.XXXX8103). This work was supported, in part, by NIH grants AG-067419, NS-42867, and NS-092988 to WDH and CCS. All aspects of this research conformed to existing US and NIH federal policies on the ethical use of chimpanzees in research. We thank Hannah Gerbeth for her support during the FreeSurfer white matter segmentation.

## Data and Code Availability

The probabilistic chimpanzee atlases of BA44 and 45, alongside the respective processing scripts used to generate them are publicly available for download in our repository: https://github.com/gagdiez/chimpanzee-broca.

